# Wheat MYB transcription factor TaMYB83-7B regulates seed dormancy by influencing the balance between abscisic acid and gibberellin

**DOI:** 10.64898/2026.05.19.726193

**Authors:** Qishi Zhuang, Shujun Cao, Litian Zhang, Huanfeng Wang, Weizhen Li, Ziwei Wang, Guanju Zhu, Wenbo Lu, Chaoxu He, Wei Gao, Can Chen, Chuanxi Ma, Haiping Zhang, Cheng Chang

## Abstract

In wheat, weak seed dormancy (SD) is related to an increased tendency for pre-harvest sprouting (PHS), which reduces yield and quality. However, the molecular mechanism underlying SD remains elusive. Here, we identified a wheat R2R3-MYB transcription factor (*TaMYB83-7B*) related to SD. Expression analysis showed that *TaMYB83-7B* was highly expressed in wheat seeds, and was more highly expressed in strong-dormancy varieties than in weak-dormancy varieties. Sequence and association analysis indicated that T/C mutations at −907 bp and −1133 bp in the *TaMYB83-7B* promoter were significantly associated with wheat SD, with C at both sites related to strong dormancy. Dual-luciferase reporter assays demonstrated that the transcriptional activity of the *TaMYB83-7B* promoter was significantly higher in strong-dormancy varieties than in weak-dormancy varieties. Further analyses indicated that TaMYB83-7B functions as a transcriptional inhibitor. Germination experiments revealed that overexpression of *TaMYB83-7B* significantly enhanced SD, while its loss-of-function reduced SD. Finally, *TaMYB83-7B* was found to regulate SD by influencing the balance between abscisic acid (ABA) and gibberellin (GA) in wheat seeds. Overall, the results of this study enhance our understanding of the complex regulatory mechanism underlying SD, and provide gene targets and molecular markers for the genetic improvement of PHS resistance in wheat.

## Introduction

Wheat (*Triticum aestivum* L.) is a global staple food crop but is highly susceptible to pre-harvest sprouting (PHS), in which wheat spikes germinate prior to harvest in response to rainfall or high humidity (Park *et al*., 2022). PHS triggers a series of physiological and biochemical reactions within wheat grains, leading to the hydrolysis of stored substances and adversely affecting yield and quality (Abhinandan *et al*., 2018; Hura, 2020). The strength of seed dormancy (SD) largely determines PHS resistance (Gubler *et al*., 2005). It is therefore critical to explore the molecular regulatory mechanism underlying SD in order to develop PHS resistant wheat varieties.

SD is a quantitative trait influenced by multiple genes, including *TaMFT* (*TaPHS1*) (Nakamura *et al*., 2011; Liu *et al*., 2013), *TaMKK3* (Torada *et al*., 2016), *TaJAZ1* (Ju *et al*., 2019), *TaQsd1* (Wei *et al*., 2019), *TaMyb10-D* (Lang *et al*., 2021), *TaSD6* (Xu *et al*., 2022), *TaGATA1* (Wei *et al*., 2023), *TaPI4K-2A* (Tai *et al*., 2024), *TaABI5-A4* (Han *et al*., 2024), *TaSRO1* (Liu *et al*., 2024), *TaPP2C-a6* (Zhang *et al*., 2024), *TaPKL* (Bai *et al*., 2025), *TaRBP-4A* (Guo *et al*., 2025), *GSK3* (Dong *et al*., 2026), *TaCNGC-2A* (Tian *et al*., 2026), *TaGASR25* (Cheng *et al*., 2026), and *TaCYP94-A1* (Zhang *et al*., 2026). These genes encode kinases, phosphatases, transcription factors (TFs), and calcium channels that regulate SD through crosstalk with phytohormones such as abscisic acid (ABA), gibberellic acid (GA), and jasmonic acid (JA). For example, TaJAZ1 negatively regulates ABA signaling and promotes seed germination by inhibiting the transcriptional activation of the ABA-responsive factor TaABI5 (Ju *et al*., 2019). The R2R3-MYB TF Myb10-D binds to the SMRE element in the *NCED* promoter, increasing ABA accumulating and conferring PHS resistance (Lang *et al*., 2021). Similarly, the TF TaGATA1 binds to the GATA motif in the *TaABI5* promoter, enhancing both ABA signaling and SD (Wei *et al*., 2023). On the other hand, TaSRO1 physically interacts with the core ABA signaling TF TaVP1, attenuating ABA signaling and reducing SD (Liu *et al*., 2024). TaPP2C-a6 physically interacts with the core dormancy regulatory proteins TaDOG1L1 and TaDOG1L4 to negatively regulate ABA signaling, thereby reducing resistance to PHS (Zhang *et al*., 2025). In wheat, PKL improves PHS resistance by modulating ABA signaling (Bai *et al*., 2025), and GSK3 enhances SD and PHS resistance by phosphorylating DOG1L4. Simultaneously, GSK3 interacts with ABI5, upregulating ABA synthesis while downregulating GA and brassinosteroid (BR) levels (Dong *et al*., 2026). Finally, TaCNGC-2A negatively regulates SD and PHS resistance by mediating Ca^2+^ and phytohormone signaling (Tian *et al*., 2026).

One of the largest TF families in plants, MYB TFs are classified into four subfamilies based on the number and position of tryptophan repeats within the conserved MYB domain: R1R2R3-MYB (3R-MYB), R2R3-MYB (2R-MYB), 4R-MYB, and MYB-like proteins. Among these, R2R3-MYB TFs constitute the largest subfamily, characterized by an N-terminal MYB DNA-binding domain and a C-terminal activation or repression domain (Dubos *et al*., 2010; Stracke *et al*., 2001; Yao *et al*., 2020). R2R3-MYB TFs are involved in the response to abiotic stresses such as drought, salinity, and low temperature (Yan *et al*., 2025; Zhou *et al*., 2025); the response to biotic stresses such as pathogens and pests (Li *et al*., 2025; Luo *et al*., 2025); and the biosynthesis of pigments and cells walls, as well as the development of seed coat color, seed size, and pollen (Cai *et al*., 2025; Luo *et al*., 2025; Zhang *et al*., 2025; Zhao *et al*., 2025). Several R2R3-MYB TFs have been found to regulate SD and germination. For example, ZmMYB59 negatively regulates seed germination in tobacco and rice by inhibiting the biosynthesis of GA_1_ and cytokinin (CK) while promoting the accumulation of ABA (Zhai *et al*., 2020). NtMYB330 enhances seed coat-imposed dormancy in tobacco by forming a specific MBW complex that regulates the biosynthesis of proanthocyanidins in the seed coat. In addition, NtMYB330 regulates the expression of ABA and GA signaling-related genes, thereby influencing seed germination (Zhao *et al*., 2022). In wheat, TaMyb10-D has been confirmed to enhance PHS resistance by directly activating the transcription of the *NCED* (9-cisepoxycarotenoid dioxygenase) gene, thereby promoting ABA biosynthesis (Lang *et al*., 2021).

Although many genes related to SD have been identified, the molecular mechanism controlling this trait in wheat remains unclear. Furthermore, relatively little is known regarding the importance of R2R3-MYB TFs in regulating wheat SD and PHS resistance. In this study, we identified the R2R3-MYB TF gene *TaMYB83-7B* (*TraesCS7B02G112400*) as well as two SD-associated single nucleotide polymorphism (SNP) mutations within its promoter. Significant differences in *TaMYB83-7B* promoter activity were observed between strong- and weak-dormancy varieties, and TaMYB83-7B was found to possess transcriptional repression activity. Further analyses suggested that *TaMYB83-7B* positively regulates SD. The results of this study provide novel insights into the molecular mechanisms underlying SD as well as genetic resources for enhancing PHS resistance in wheat.

## Materials and Methods

### Plant materials and growth conditions

Six wheat cultivars with contrasting dormancy levels were used as experimental materials to investigate the role of *TaMYB83-7B* in SD and germination. The weak-dormancy (PHS sensitive) cultivars included Jing 411 (J411), Zhongyou 9507 (ZY9507), and Zhoumai 22 (ZM22). The strong-dormancy (PHS resistant) cultivars included Hongmangchun 21 (HMC21), Yangxiaomai (YXM), and Yangmai 16 (YM16). Plants were cultivated at the Anhui Agricultural University Dayangdian Experimental Station in Hefei, China (31°93′N, 117°20′E) using standard agronomic practices. To investigate *TaMYB83-7B* expression, grain samples were collected at 25, 30, and 35 days post-anthesis (DPA; representing the dormancy establishment process) and at 5, 15, and 30 days after harvest (DAH; representing the dormancy release process).

Thirty healthy, fully mature seeds per variety were placed in sterile Petri dishes double-lined with sterile filter paper moistened with sterile distilled water. Seeds were incubated at 22°C (day)/20°C (night) under a 16-h light/8-h dark photoperiod with 70% relative humidity. Seed samples were collected following 4, 6, and 10 h of imbibition, immediately frozen in liquid nitrogen, and stored at −80°C for subsequent RNA extraction. Three biological replicates were prepared for each time point.

Associations between *TaMYB83-7B* and SD were investigated using 192 wheat varieties (192WVs). Germination index (GI) values were calculated at 5, 15, and 30 DAH during the 2014-2015, 2015-2016, and 2016-2017 cropping seasons, and were designated as 15GI-5DAH, 15GI-15DAH, 16GI-5DAH, 16GI-15DAH, 16GI-30DAH, 17GI-5DAH, and 17GI-15DAH, respectively.

The ethyl methane sulfonate (EMS)-induced mutant *tamby83-7b* (chromosome 7B mutation; ID 7B_131642395_E) was provided by MolBreeding (https://jing411.molbreeding.com/#/query). To minimize the effect of background mutations, this mutant was backcrossed three times to the J411 population, followed by selfing to produce BCLFL progeny for phenotypic and molecular analyses. The *tamyb83-7b* mutant was cultivated at the Anhui Agricultural University Dayangdian Experimental Station in Hefei, China (31°93′N, 117°20′E).

*TaMYB83-7B* overexpression (OE) lines were constructed in the Fielder background. OE plants were grown in a growth chamber at 23°C ± 2°C under a 16-hour light/8-hour dark photoperiod and 70% relative humidity. Only homozygous T_2_ transgenic plants were used for subsequent analyses.

The *TaMYB83-7B* coding sequence (CDS) was amplified from J411 cDNA and cloned into the pBWA(V)HS-GUS plant expression vector using the *Bsa* I/*Eco31* I restriction sites. Subsequently, the recombinant vector was introduced into *Arabidopsis thaliana* Col-0 and Nipponbare (Nip) rice using *Agrobacterium*-mediated flower immersion (Clough and Bent, 1998) and rice callus transformation (Slamet-Loedin *et al*., 2014), respectively.

## Germination assays

The germination rates of wheat, *A. thaliana*, and rice seeds were evaluated. The GP was calculated as the ratio of the number of germinated seeds to the total number of seeds according to Xu *et al*. (2022).

Wheat seeds collected at 35 DPA were surface sterilized with 75% alcohol for 15 min and subsequently washed five times (3 min each) with sterile water. Sterilized seeds were sown in circular glass Petri dishes with the ventral groove facing downward. The germination percentage (GP) was determined using an incubator set at 22°C ± 2°C under a 16-hour light/8-hour dark photoperiod and 70% relative humidity. Seed germination was defined as the point at which the wheat radicle successfully pierced the seed coat.

Thirty rice seeds were treated with 0.1% (w/v) sodium hypochlorite for 15 min followed by five rinses (3 min each) with deionized water. Subsequently, the seeds were placed on moist filter paper and incubated at 28°C (light)/25°C (dark) with 70% relative humidity. Germination was defined as radicle length ≥ 1 mm (Li *et al*., 2020).

*A. thaliana* seeds were surface-sterilized in a 75% ethanol solution for 10 min followed by five rinses with deionized water (3 min each). Subsequently, the seeds were evenly sown on moist germination paper and incubated at 23°C (light)/20°C (dark) with 60% relative humidity. Germination was defined as radicle protrusion through the seed coat and was scored daily until all seeds had germinated (Cao *et al*., 2020).

To simulate field conditions conducive to PHS, wheat spikes harvested at approximately 35 DPA (physiological maturity) and freshly harvested rice panicles were immersed in a 0.1% sodium hypochlorite solution for 15 min, rinsed three times with tap water (5 min each), and vertically mounted on a stainless steel rack equipped with spray nozzles, where they were sprayed every 8 hr.

### Phylogenetic analysis

Protein sequences homologous to TaMYB83-7B were identified by BLAST search against NCBI (https://www.ncbi.nlm.nih.gov) and Ensembl Plants (http://plants.ensembl.org). Multiple sequence alignments were performed using DNAMAN v8. A neighbor-joining phylogenetic tree was constructed with MEGA11, and branch support was evaluated by 1,000 bootstrap replicates (Tamura *et al*., 2021).

### RNA extraction and RT-qPCR analysis

Total RNA was extracted from wheat, rice, and *A. thaliana* seeds at different developmental stages using a plant RNA extraction kit (DP220706, TiGen Bio, Beijing, China). First-strand cDNA was synthesized in a 40 µL reaction system using a FastKing RT Kit containing gDNase (KR116, TiGen Bio). Reference genes included *TaActin1* for wheat (Paolacci *et al*., 2009), *OsActin1* for rice (Wang and Bhullar, 2021), and *AtActin7* for *A. thaliana* (Jin *et al*., 2019). RT-qPCR was performed on a CFX96 real-time system (Bio-Rad, USA) in a 20 µL reaction volume. Relative expression levels were calculated using the 2^DΔΔCt^ method. Data processing and visualization were performed with Excel and GraphPad Prism v8.

### Gene cloning and molecular marker development

The full-length *TaMYB83-7B* sequence was retrieved from the IWGSC RefSeq v2.0 database. Gene-specific primers (Supplementary Table S1) were designed to amplify the gene from J411, ZY9507, ZM22, HMC21, YXM, and YM16 wheat. PCR was performed using TransFast Taq DNA Polymerase (TransGen Biotech, Beijing, China) in a 20 µL reaction volume. The amplicons were separated on 1.8% agarose gels, and the target bands were purified (TIANgel Purification Kit, TIANGEN) and sequenced. Sequence alignment (DNAMAN) was performed to identify SNPs in the promoter region as well as their flanking restriction sites. Four cleaved amplified polymorphic sequence (CAPS) markers were developed based on the identified SNPs: TaMYB83-7B-736, TaMYB83-7B-907, TaMYB83-7B-1133, and TaMYB83-7B-1321 (Shavrukov, 2016).

### Dual-luciferase reporter assay

The transcriptional activation activity of TaMYB83-7B was evaluated via dual-luciferase (dual-LUC) reported assay (Hellens *et al*., 2005). The *TaMYB83-7B* CDS was inserted into pGreenII-62SK-GAL4-BD under the control of the CaMV 35S promoter. GAL4-BD served as a negative control and GAL4-BD-VP16 served as a positive control. The pGreenII-0800-LUC-GAL4-TATA reporter construct contains GAL4 binding sites, and renilla luciferase (REN) was used as the internal reference. The plasmids were transformed into *Agrobacterium* GV3101 (pSoup-19), which was applied to the undersides of *Nicotiana benthamiana* leaves. LUC signals were observed using a Tanon 5200 chemiluminescence system. LUC and REN activities were quantified using a Dual-Luciferase Reporter Assay Kit (DL101-01, Vazyme Biotech, Nanjing) in order to calculate the LUC/REN ratio.

To verify the effects of different *TaMYB83-7B* promoter alleles on transcriptional activity, promoter fragments from HMC21 (TaMYB83-7B^HMC21Pro^) and J411 (TaMYB83-7B^J411Pro^) were cloned into pGreenII-0800-LUC. An empty LUC vector served as a negative control. All other assay conditions were as described above.

### Quantification of GA and ABA contents and **α**-amylase activity

*TaMYB83-7B-OE#7* and Fielder seeds were harvested at 35 DPA (physiological maturity), placed in petri dishes containing 8 mL of distilled water, and germinated in a constant-temperature incubator at 22 ± 2°C. Samples were collected following 0, 12, and 24 h of imbibition. GA and ABA contents were measured with plant hormone-specific enzyme-linked immunosorbent assay (ELISA) kits (BY-JZF0146 and BY-JZF0147, BYabscience Biotech, Shanghai, China). Absorbance (450 nm) was quantified using a SPARK multifunctional microplate reader (TECAN, Switzerland), and hormone contents were calculated from standard curves. The activity of α-amylase was assayed using a commercial colorimetric kit (BC0615, Solarbio, Beijing, China). All assays included three biological replicates, each comprising three technical replicates.

### Subcellular localization analysis

To determine the subcellular localization of TaMYB83-7B, the *TaMYB83-7B* CDS was cloned into p1305-35S-GFP to generate the 35S-TaMYB83-7B-GFP fusion construct. 35S-GFP served as an empty control. The nuclear marker plasmid mCherry-H2B (pABR113) under control of the CaMV 35S promoter was co-transformed to delineate nuclear boundaries. Rice mesophyll protoplasts were isolated and transformed via PEG-mediated transfection. After incubation in the dark at 22°C for 16 h, GFP and mCherry signals were observed using a Nikon C2-ER confocal laser scanning microscope (Nikon, Tokyo, Japan) (Yoo *et al*., 2007).

## Results

### Identification and expression analysis of TaMYB83-7B

To investigate the role of the *MYB* gene family in wheat SD, we compared the transcriptomes of strong- (HMC21) and weak-dormancy (J411) seeds following 4, 6, and 10 h of imbibition. All J411 seeds germinated by 10 h of imbibition, whereas HMC21 seeds did not. Four *MYB* genes exhibited differential expression between J411 and HMC21 seeds following 10 h of imbibition: *TraesCS3A02G414100*, *TraesCS1B02G236900*, *TraesCS7B02G112400*, and *TraesCS5B02G416500* (Fig. 1A. Supplementary Fig. S1A). Subsequent RT-qPCR analyses confirmed these expression trends (Fig. 1B).

**Fig. 1.**
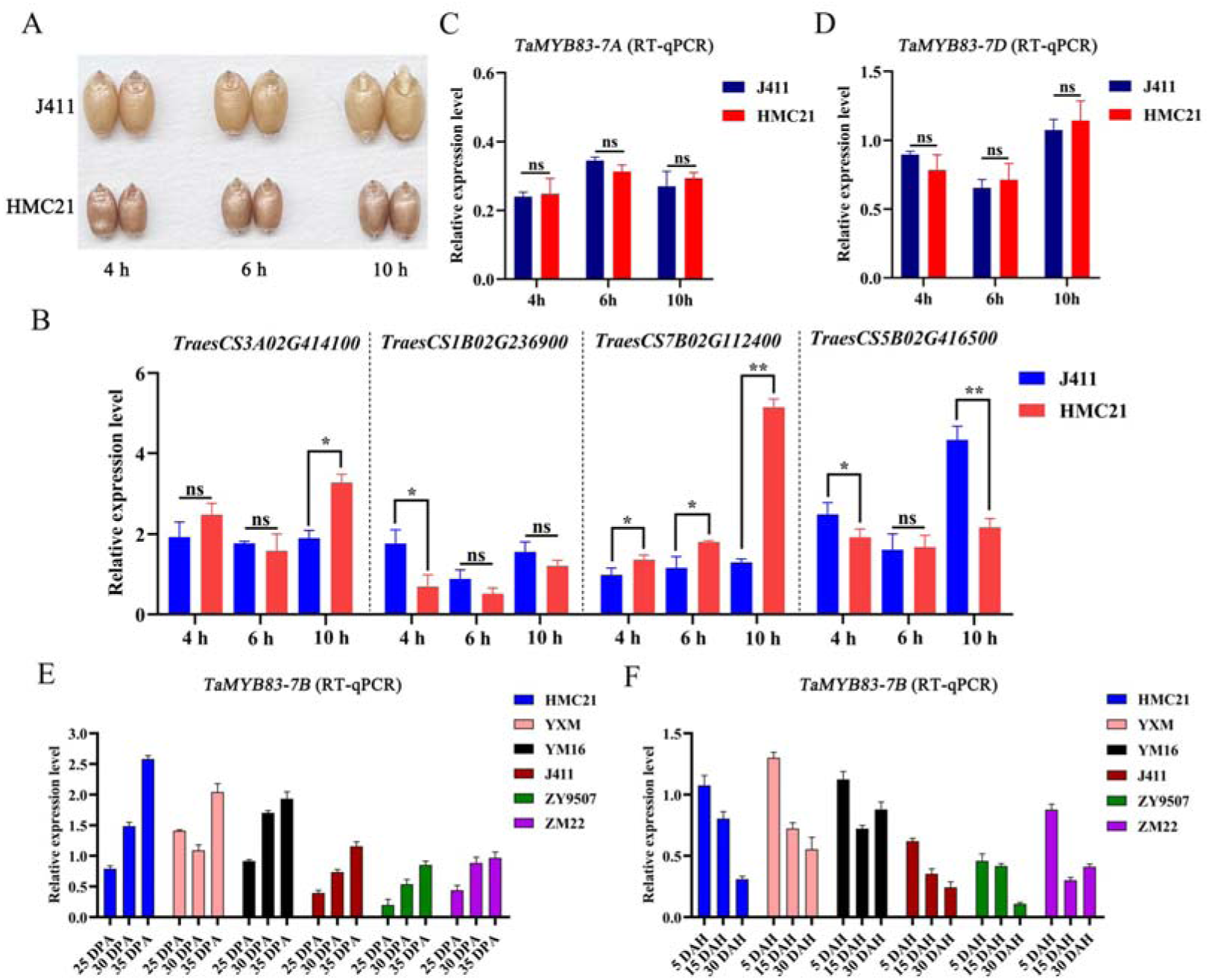
Identification and expression analysis of *TaMYB83-7B*. **A.** J411 and HMC21 wheat seeds during germination. **B.** RT-qPCR analysis of *TraesCS3A02G414100*, *TraesCS1B02G236900*, *TraesCS7B02G112400,*and *TraesCS5B02G416500* expression in J411 and HMC21 seeds imbibed for 4, 6, and 10 h. **C-D.** RT-qPCR analysis of *TaMYB83-7A* (**C**) and *TaMYB83-7D* (**D**) expression in J411 and HMC21 seeds imbibed for 4, 6, and 10 h. **E.** Expression of *TaMYB83-7B* in HMC21, YXM, YM16, J411, ZY9507, and ZM22 wheat seeds at 25, 30, and 35 days post-anthesis (DPA). **F.** Expression of *TaMYB83-7B* in HMC21, YXM, YM16, J411, ZY9507, and ZM22 wheat seeds at 5, 15, and 30 days after harvest (DAH). Data are shown as means ± SD (n = 3). Statistical significance was assessed using Student’s *t*-test (\*\**P* < 0.01). ns, “not significant”.

The expression patterns of these four genes in two weak-dormancy varieties (ZY9507 and ZM22) and two strong-dormancy varieties (YXM and YM16) were analyzed following 4, 6, and 10 h of imbibition. Only *TraesCS7B02G112400* exhibited consistently significantly different expression between weak- and strong-dormancy varieties (Supplementary Fig. S1B). RT-qPCR analysis showed that *TraesCS7B02G112400* was highly expressed specifically in seeds (Supplementary Fig. S1C-F). Together, these results suggest that *TraesCS7B02G112400* (designated *TaMYB83-7B*) may be a candidate gene controlling SD.

We next analyzed the expression patterns of the *TaMYB83-7B* homologs *TaMYB83-7A* and *TaMYB83-7D*. Following 4, 6, or 10 h of imbibition, no significant differences in *TaMYB83-7A* and *TaMYB83-7D* expression were observed between J411 and HMC21 seeds (Supplementary Fig. S2). Subsequent RT-qPCR analyses confirmed these expression trends (Fig. 1C-D). Therefore, only *TaMYB83-7B* was selected for further analysis.

*TaMYB83-7B* expression was evaluated in six wheat cultivars with contrasting dormancy levels at 25, 30, and 35 DPA (the dormancy establishment process) and 5, 15, and 30 DAH (the dormancy release process). *TaMYB83-7B* expression was higher during dormancy establishment but decreased significantly during dormancy release. In addition, *TaMYB83-7B* expression was consistently higher in strong-dormancy cultivars (HMC21, YXM, and YM16) than in weak-dormancy cultivars (J411, ZY9507, and ZM22) (Fig. 1E-F). These results suggest that *TaMYB83-7B* may particulate in dormancy establishment and release.

### Subcellular localization, phylogenetic analysis, and transcriptional activity of TaMYB83-7B

We observed that the green fluorescence signal of 35S:TaMYB83-GFP was specifically localized to the nucleus and overlapped completely with the red fluorescence signal of the nuclear marker protein mCherry (Fig. 2A). These results indicate that TaMYB83 is a nuclear-localized protein.

**Fig. 2.**
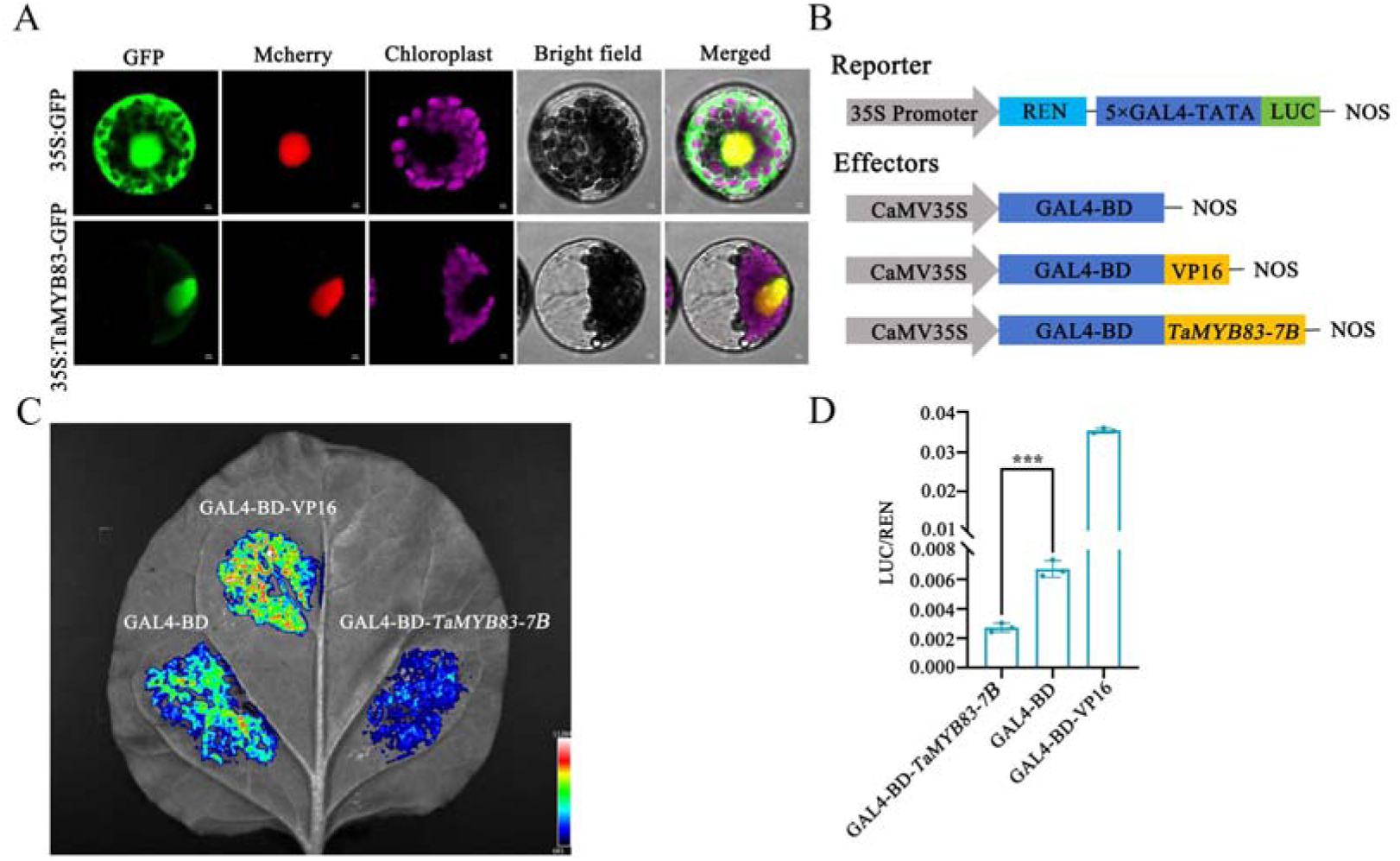
Subcellular localization and transcriptional activation analysis of TaMYB83-7B. **A.** Subcellular localization of TaMYB83-7B in wheat protoplasts. Scale bars = 10 µm. **B.** Reporter genes and effector elements used in the transcriptional activity assay. **C-D.** Transcriptional activation analysis of TaMYB83-7B. Dual-luciferase reporter assay revealing TaMYB83-7B to be a transcriptional repressor. Data are shown as means ± SD (n = 3). Statistical significance was assessed using Student’s *t*-test (\*\*\**P* < 0.001).

Additionally, a phylogenetic tree was constructed using 15 rice and 11 *A. thaliana* MYB protein sequences. The TaMYB83-7B protein was classified into a distinct clade and shared 47.35% homology with AtMYB20 (AT1G66230), a member of the *A. thaliana* R2R3-MYB TF family (Supplementary Fig. S3), suggesting that TaMYB83-7B is a R2R3-MYB-type TF.

To determine whether TaMYB83-7B possesses transcriptional activation activity, we constructed a GAL4-based dual-LUC reporter system and performed transient expression in tobacco leaves (Fig. 2B). Chemiluminescence imaging revealed that TaMYB83-7B expression resulted in significantly lower LUC activity compared to the empty vector control, indicating that TaMYB83-7B exhibits transcriptional repressor activity in plant cells (Fig. 2C). In addition, the observed LUC/REN ratio is consistent with the results of chemiluminescence imaging analysis (Fig. 2D). These results suggest that TaMYB83-7B functions as a transcriptional repressor.

### TaMYB83-7B promoter sequence variations are associated with SD

Based on the Chinese Spring wheat genome, we cloned the *TaMYB83-7B* genomic sequence from six wheat varieties with contrasting dormancy levels (HMC21, YXM, YM16, J411, ZY9507, and ZM22). The *TaMYB83-7B* gene spans 2922 bp, including a 1500 bp promoter, a 174 bp 5’ UTR, three exons, two introns, and a 252 bp 3’ UTR. The *TaMYB83-7B* CDS is 795 bp in length and encodes 264 amino acids. Four SNPs were identified in the *TaMYB83-7B* promoter between weak- and strong-dormancy varieties, while no variants were observed in the coding region. These four SNPs resulted in changes to transcription factor binding sites within the *TaMYB83-7B* promoter. In J411 relative to HMC21, the G-to-A mutation at −736 bp resulted in four additional TCR/CPP binding sites, the C-to-T mutations at −907 bp and v1133 bp resulted in seven additional bHLH binding sites, and the A-to-G mutation at −1321 bp resulted in the loss of one Dof binding site and the addition of one AP2/ERF binding site (Fig. 3A).

**Fig. 3.**
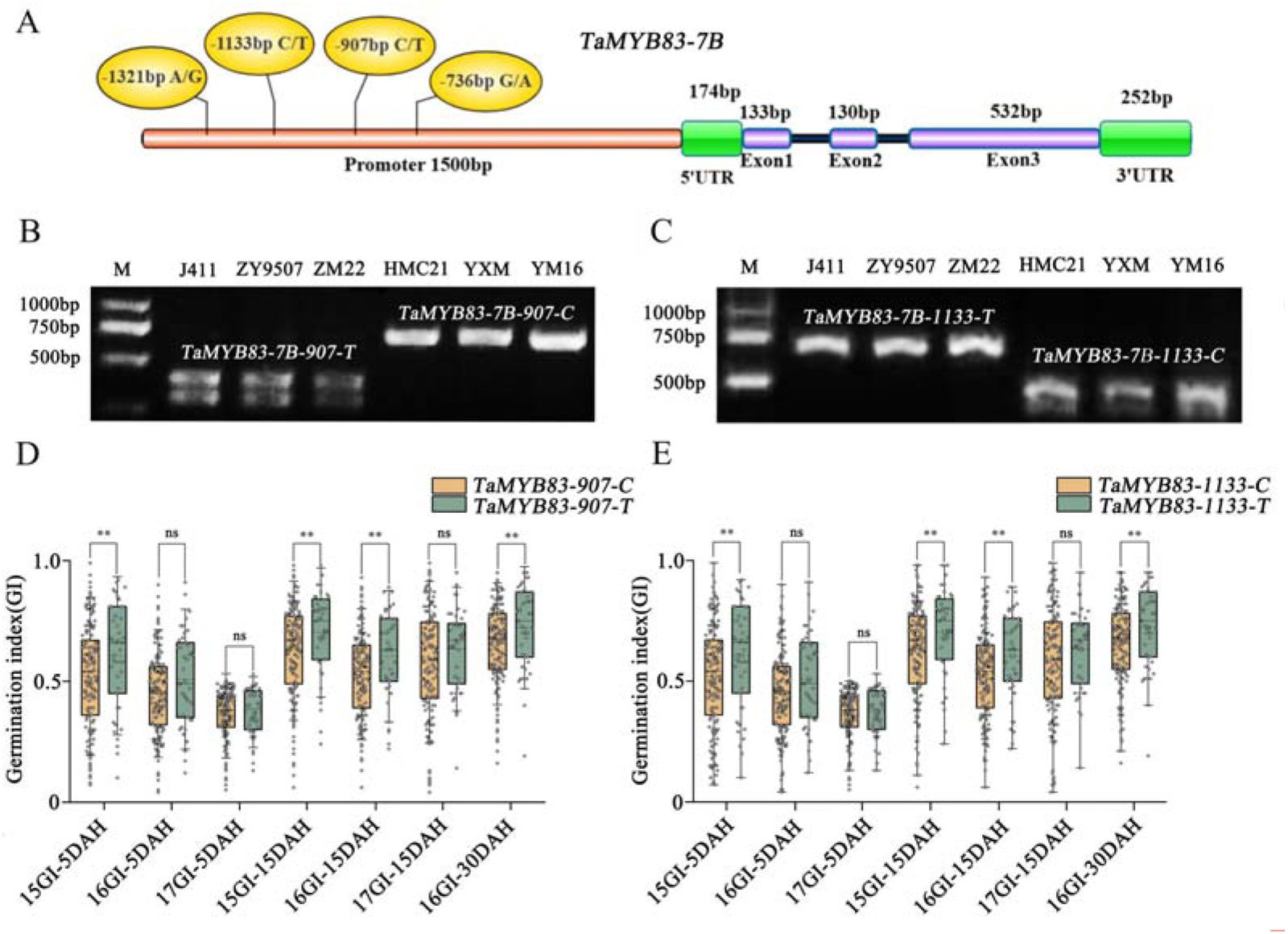
Sequence variations in the *TaMYB83-7B* promoter are associated with seed dormancy. **A.** *TaMYB83-7B* gene structure. **B-C.** Detection of two CAPS markers (TaMYB83-7B-907 and TaMYB83-7B-1133) for *TaMYB83-7B* in J411, ZY9507, ZM22, HMC21, YXM, and YM16 wheat using *BssH* II and *Nsi* I on 1.8% agarose gels. M represents the 2 K DNA marker. **D.** Significant differences were observed in germination index (GI) values among 192 wheat varieties (192WVs) with the two allelic variations of *TaMYB83-7B-907* (*TaMYB83-7B-907-T* and *TaMYB83-7B-907-C*). **E.** Significant differences were observed in GI values among 192WVs with the two allelic variations of *TaMYB83-7B-1133* (*TaMYB83-7B-1133-T* and *TaMYB83-7B-1133-C*). 15GI-5DAH, 16GI-5DAH, 17GI-5DAH, 15GI-15DAH, 16GI-15DAH, 17GI-15DAH, and 16GI-30DAH represent GI values measured 5, 15, and 30 days after harvest (DAH) in 2015, 2016, and 2017, respectively. Data are shown as means ± SD (n = 3). Statistical significance was assessed using Student’s *t*-test (\*\**P* < 0.01). ns, “not significant”.

To assess the effects of these four SNPs on SD, we developed four CAPS markers (TaMYB83-7B-736, TaMYB83-7B-907, TaMYB83-7B-1133, and TaMYB83-7B-1321) to genotype 192WVs with different dormancy levels (Supplementary Table S2). The genotypes detected by TaMYB83-7B-907 and TaMYB83-7B-1133 were identical (Fig. 3B-C. Supplementary Fig. S4A-B).

Mann-Whitney U test analysis revealed significant differences in GI between varieties carrying the two alleles of TaMYB83-7B-907 (*TaMYB83-7B-907-T* and *TaMYB83-7B-907-C*) and TaMYB83-7B-1133 (*TaMYB83-7B-1133-T* and *TaMYB83-7B-1133-C*), whereas no significant differences were observed between those carrying the two alleles of TaMYB83-7B-736 (*TaMYB83-7B-736-A* and *TaMYB83-7B-736-G*) and TaMYB83-7B-1321 (*TaMYB83-7B-1321-G* and *TaMYB83-7B-1321-A*) (Supplementary Table S3). *TaMYB83-7B-907-C* and *TaMYB83-7B-1133-C* were associated with lower GI (strong dormancy), while *TaMYB83-7B-907-T* and *TaMYB83-7B-1133-T* were associated with higher GI (weak dormancy) (Fig. 3D-E. Supplementary Fig. S4C-D). These results suggest that the T/C mutations at −907 bp and −1133 bp are significantly associated with SD.

### Two SNPs at −907 bp and −1133 bp affect TaMYB83-7B promoter activity

To explore whether the four identified SNPs affect the transcriptional activity of TaMYB83-7B, we performed dual-LUC reporter assays in tobacco. The transcriptional activity of TaMYB83-7B^HMC21Pro^ (HMC21 promoter) was significantly higher than that of TaMYB83-7B^J411Pro^ (J411 promoter) (Fig. 4A). To determine which SNP is responsible for the difference in transcriptional activity between TaMYB83-7B^HMC21Pro^ and TaMYB83-7B^J411Pro^, we conducted promoter site-directed mutagenesis. Specifically, using the HMC21-type promoter sequence as a template, we converted the four SNPs into J411-type genotypes. We constructed promoter reporter vectors containing a mutation at only the −736 bp site (TaMYB83-7B^M1Pro^), a mutation at only the −1321 bp site (TaMYB83-7B^M3Pro^), and a double mutation simultaneously at both the −907 bp and −1133 bp sites (TaMYB83-7B^M2Pro^) (Fig. 4B). The SNPs at −907 bp and −1133 bp significantly affected TaMYB83-7B promoter activity, while the SNPs at −736 bp and −1321 bp did not (Fig. 4C).

**Fig. 4.**
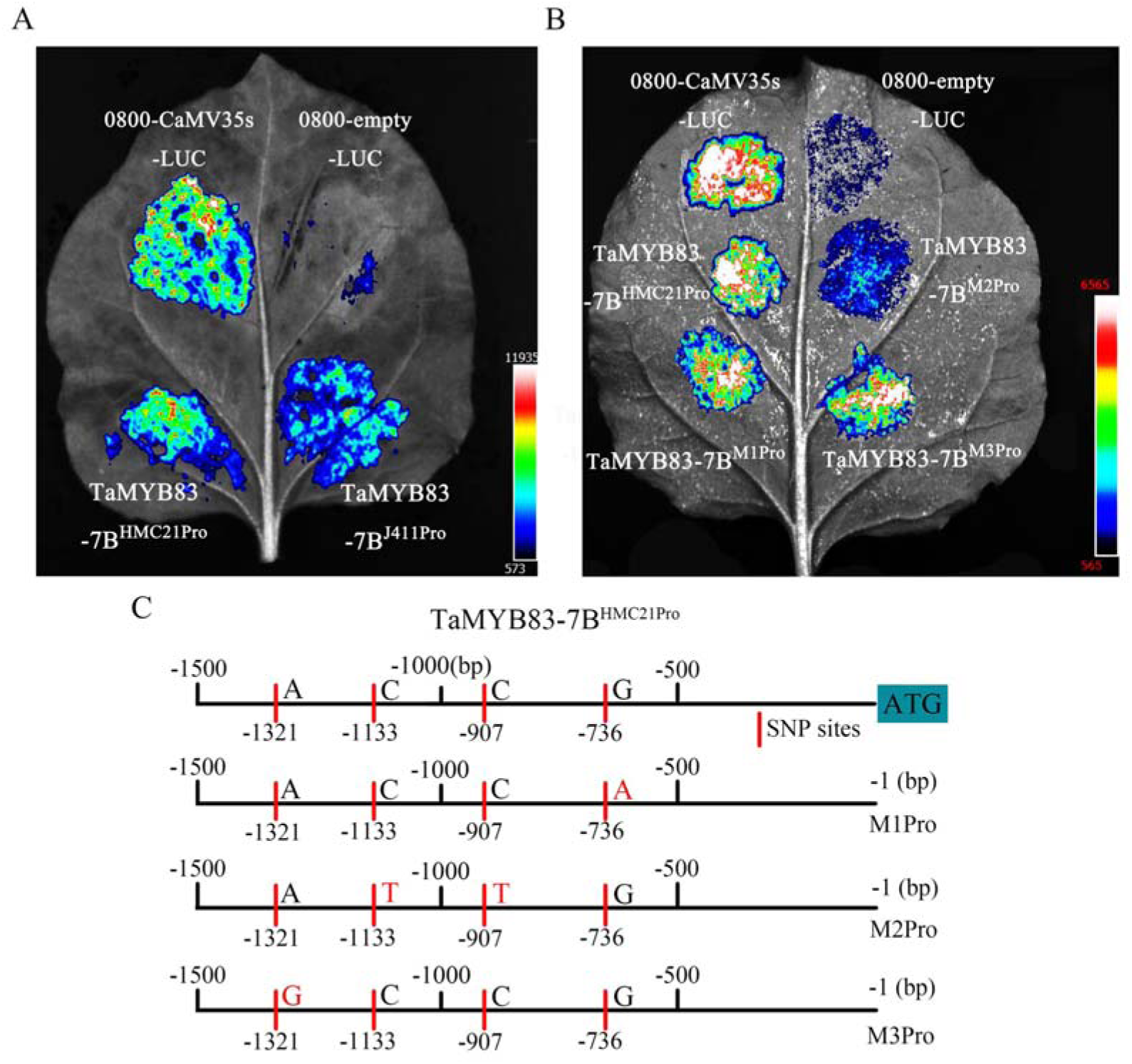
Two single nucleotide polymorphisms (SNPs) at −907 bp and −1133 bp affect *TaMYB83-7B* promoter activity. **A.** Analysis of *TaMYB83-7B* promoter activity. Dual-luciferase reporter assays demonstrating that the TaMYB83-7B^HMC21Pro^ promoter exhibits significantly higher activity than the TaMYB83-7B^J411Pro^ promoter. 0800-CaMV35s-LUC served as the positive control while 0800-empty vector-LUC served as the negative control. **B-C.** Using the TaMYB83-7B^HMC21Pro^ promoter sequence as a template, site-directed mutagenesis was performed at −736, −907, −1133, and −1321 Lbp. Mutation at −736L bp yielded TaMYB83-7B^M1Pro^, simultaneous mutations at −907L bp and −1133L bp yielded TaMYB83-7B^M2Pro^, and mutation at −1321Lbp yielded TaMYB83-7B^M3Pro^. Dual-luciferase reporter assays showing that the activities of the TaMYB83-7B^HMC21Pro^, TaMYB83-7B^M1Pro^, and TaMYB83-7B^M3Pro^ promoters were not significantly different from each other but were significantly higher than that of the TaMYB83-7B^M2Pro^ promoter. 0800-CaMV35s-LUC served as the positive control while the 0800-empty vector-LUC served as the negative control.

### The role of TaMYB83-7B in wheat SD

To validate the role of *TaMYB83-7B* in wheat SD, we generated *TaMYB83-7B* OE lines (*TaMYB83-7B-OE#2*, *TaMYB83-7B-OE#4*, and *TaMYB83-7B-OE#7*) in the Fielder background (Fig. 5A). Compared with Fielder (87%), the GPs of *TaMYB83-7B-OE#2* (64%), *TaMYB83-7B-OE#4* (68%), and *TaMYB83-7B-OE#7* (67%) were significantly reduced (Fig. 5B-C).

**Fig. 5.**
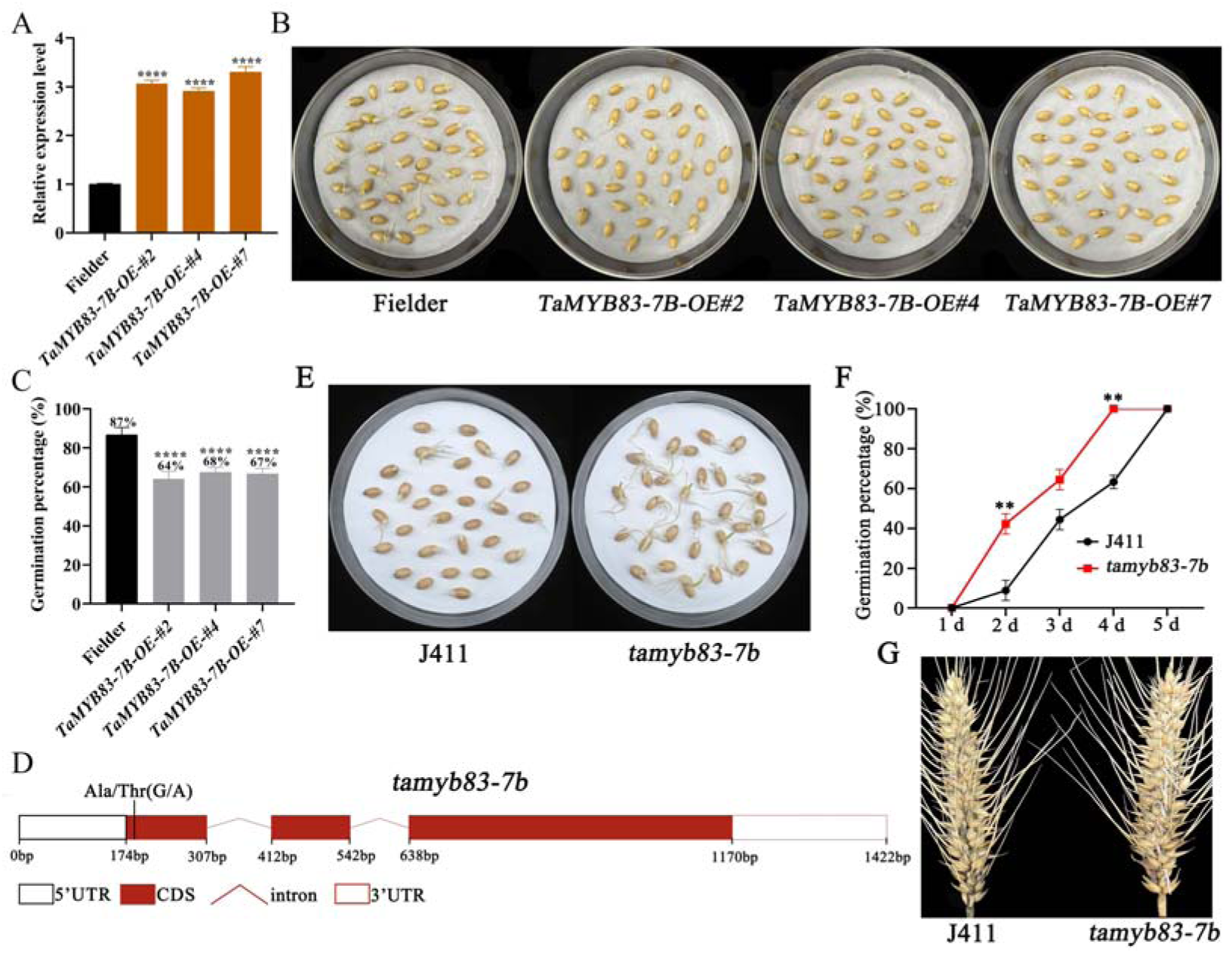
Role of *TaMYB83-7B* in wheat seed dormancy. **A.** Expression of *TaMYB83-7B* in *TaMYB83-7B-OE#2/4/7* and Fielder. **B.** Germination phenotypes of *TaMYB83-7B*L*OE#2/4/7* and Fielder seeds following 5 d of imbibition. **C.** Germination percentages (GPs) of *TaMYB83-7B*L*OE#2/4/7* and Fielder seeds. **D.** Mutation site of the wheat EMS mutant *tamyb83-7b*. **E.** Germination phenotypes of freshly harvested *tamyb83-7b* and J411 seeds imbibed for 4 d. **F.** GPs of *tamyb83-7b* and J411 seeds imbibed for 5 d. **G.** Germination phenotypes of *tamyb83-7b* and J411 spikes imbibed for 4 d. Data are shown as means ± SD (n = 3). Statistical significance was assessed using Student’s t-*t*est (\*\**P* < 0.01, \*\*\*\**P* < 0.0001).

We also obtained the EMS-induced *tamyb83-7b* mutant in the J411 background. In this mutant, a G-to-A mutation in the *TaMYB83-7B* coding region results in the substitution of the 19^th^ amino acid from nonpolar alanine (Ala) to polar threonine (Thr) within the SANT domain of the TaMYB83-7B protein (Fig. 5D). *tamyb83-7b* seeds germinated significantly faster than J411 seeds, with 40% of *tamyb83-7b* seeds germinating following 2 d of imbibition and 100% following 4 d (Fig. 5E-F). Furthermore, the germination rate of *tamyb83-7b* spikes markedly increased following 4 d of continuous spraying (Fig. 5G). These observations suggest that the *tamyb83-7b* mutant possesses significantly weaker SD than wild-type J411, and that TaMYB83-7B functions as a positive regulator of SD.

### The role of TaMYB83-7B in A. thaliana and rice SD

Many previously identified dormancy-related genes (e.g., *Vp-1/ABI3*, *MFT*, *Qsd1/AlaAT*, *Sdr*, *DOG1*, and *MKK3*) show functional conservation across plant species such as *A. thaliana*, rice, maize, barley, and wheat. To investigate the functional conservation of *TaMYB83-7B*, we first heterologously expressed this gene in *A. thaliana* Col-0. Three independent OE lines with relatively high expression levels (*TaMYB83-7B-OE-1*, *TaMYB83-7B-OE-2*, and *TaMYB83-7B-OE-3*) were selected for germination testing (Supplementary Fig. S5A). Germination was delayed in all OE lines compared with Col-0. Specifically, the average GP of the OE lines was significantly lower than that of Col-0 following 2 to 6 d of imbibition, although the GPs of both groups approached 100% by 8 d (Fig. 6A-B). These results indicate that heterologous expression of *TaMYB83-7B* enhances SD and suppresses germination in *A. thaliana*.

**Fig. 6.**
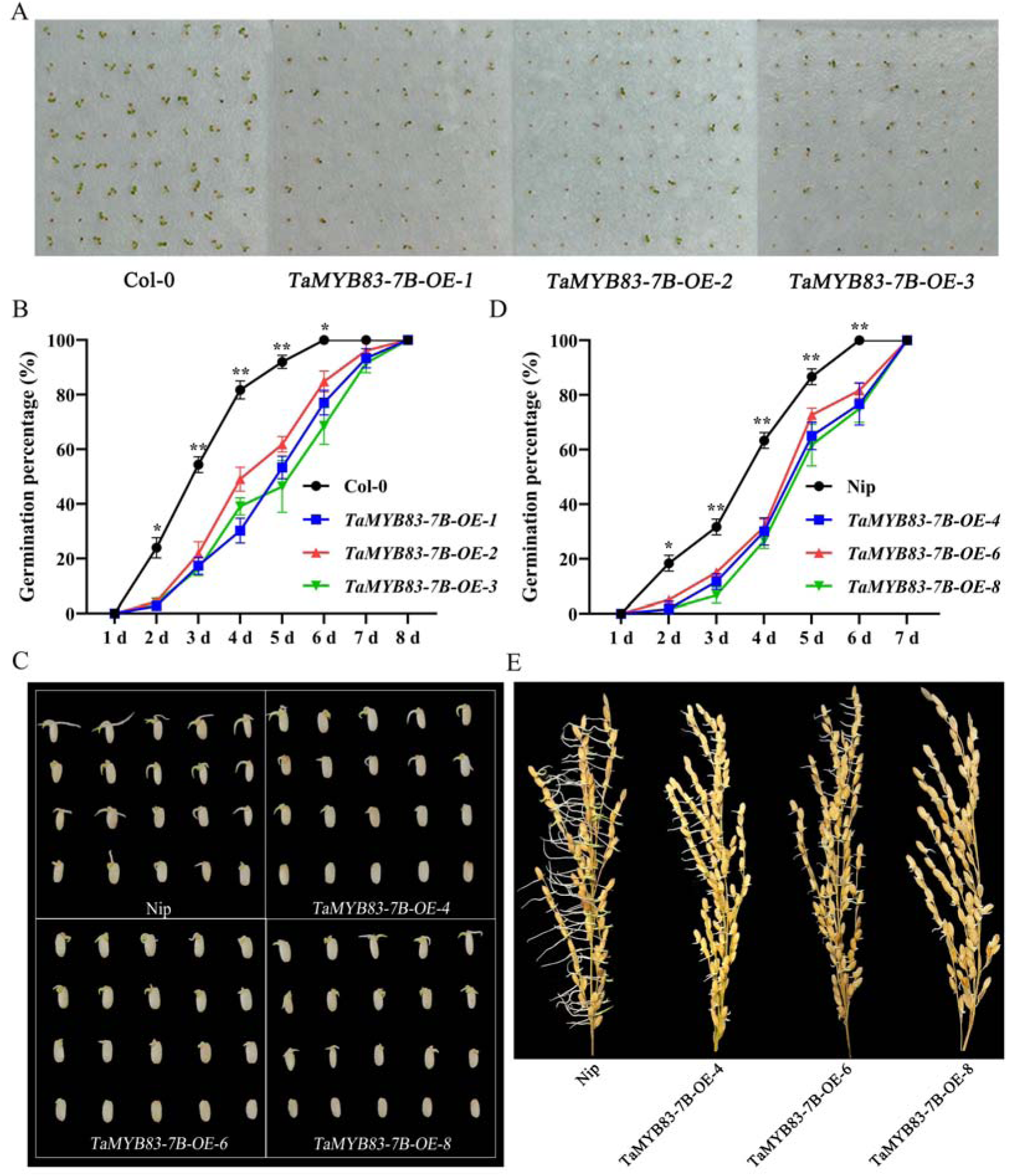
Role of *TaMYB83-7B* in *Arabidopsis thaliana* and rice seed dormancy. **A.** Germination phenotypes of *TaMYB83-7B*-overexpressing *A. thaliana* and wild-type Col-0 seeds imbibed for 5 d. **B.** Germination percentages (GP) of freshly harvested *TaMYB83-7B-OE* and Col-0 seeds. **C.** Germination phenotypes of *TaMYB83-7B-OE* rice and wild-type Nip seeds imbibed for 6 d. **D.** GPs of freshly harvested *TaMYB83-7B-OE* and Nip seeds. **E.** Germination phenotypes of *TaMYB83-7B-OE* and Nip panicles imbibed for 7 d. Data are shown as means ± SD (n = 3). Statistical significance was assessed using Student’s *t*-test (\**P* < 0.05, \*\**P* < 0.01).

We next obtained three *TaMYB83-7B* OE lines in Nip rice: *TaMYB83-7B-OE-4*, *TaMYB83-7B-OE-6*, and *TaMYB83-7B-OE-8* (Supplementary Fig. S5B). Compared with Nip, all OE lines exhibited delayed germination. The GP values of the OE lines were significantly lower than those of Nip following 2 to 6 d of imbibition, but approached 100% in both groups by 7 d (Fig. 6C-D). We also observed a significant delay in seed germination from freshly harvested mature panicles (Fig. 6E). As was the case for *A. thaliana*, heterologous expression of *TaMYB83-7B* in rice enhances SD and suppresses germination.

### TaMYB83-7B mediates SD through the ABA and GA pathways

The phytohormones ABA and GA are known to play roles in regulating SD and germination. To elucidate whether *TaMYB83-7B* regulates wheat SD and germination through crosstalk with ABA and GA, we analyzed endogenous ABA and GA levels in *TaMYB83-7B-OE#7* and Fielder seeds following 0, 12, and 24 h of imbibition. The ABA content was significantly higher in *TaMYB83-7B-OE#7* seeds than in Fielder seeds, whereas the GA content exhibited the opposite trend. Notably, the GA level in Fielder was approximately 1.4 times that in *TaMYB83-7B-OE#7* following 12 h of imbibition, leading to a significantly decreased ABA/GA ratio (Fig. 7A). The activity of α-amylase was significantly lower in *TaMYB83-7B-OE#7* seeds than in Fielder seeds at the same time point (Fig. 7B), providing a mechanistic basis for the observed delayed germination phenotype of *TaMYB83-7B-OE#7* seeds.

**Fig. 7.**
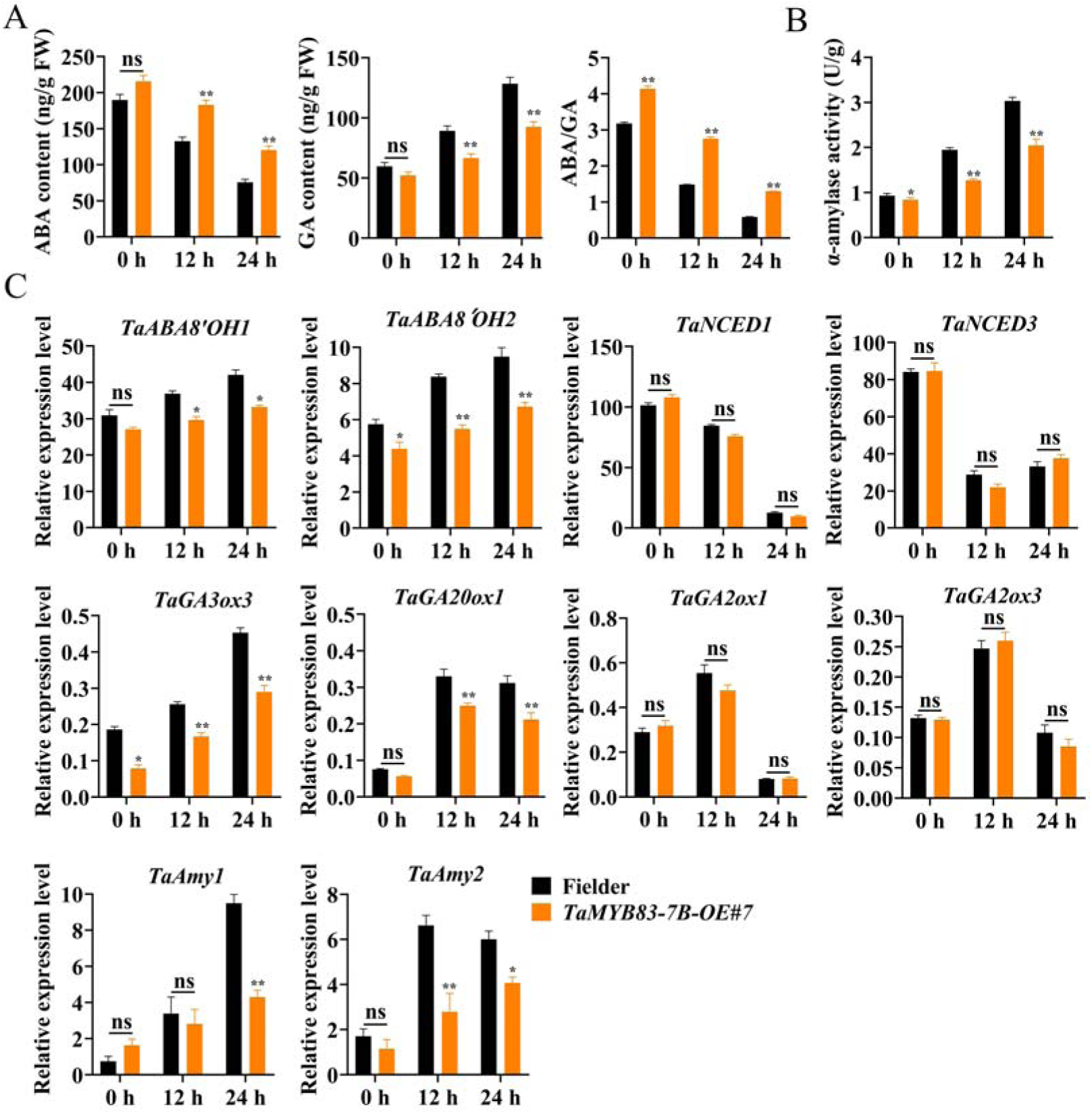
***TaMYB83-7B* mediates seed dormancy through the ABA and GA pathways. A-B.** GA and ABA contents and α-amylase activity in *TaMYB-7B-OE#7* and Fielder seeds imbibed for 0, 12, and 24 h. **C.** Relative expression levels of key genes involved in GA and ABA biosynthesis and metabolism in *TaMYB-7B-OE#7* and Fielder seeds imbibed for 0, 12, and 24 h. Data are shown as means ± SD (n = 3). Statistical significance was assessed using Student’s *t*-test (\**P* < 0.05, \*\**P* < 0.01). ns, “not significant”.

To explore how *TaMYB83-7B* influences the biosynthesis of ABA and GA at the transcriptional level, we analyzed the expression of key genes involved in ABA and GA biosynthesis and metabolism in *TaMYB83-7B-OE#7* and Fielder seeds imbibed for 0, 12, and 24 h using RT-qPCR. Compared to Fielder seeds, genes related to ABA catabolism (*TaABA8’OH1* and *TaABA8’OH2*) and GA biosynthesis (*TaGA3ox3* and *TaGA20ox1*) were significantly downregulated in *TaMYB83-7B-OE#7* seeds, especially following 12 and 24 h of imbibition. In contrast, no significant differences were observed in the expression levels of genes involved in ABA biosynthesis (*TaNCED1* and *TaNCED3*) and GA degradation (*TaGA2ox1* and *TaGA2ox3*).

Notably, the expression of the α-amylase synthesis genes *TaAmy1* and *TaAmy2* significantly decreased as imbibition time increased in *TaMYB83-7B-OE#7* seeds (Fig. 7C). Together, these findings suggest that *TaMYB83-7B* regulates the expression of multiple genes involved in ABA and GA biosynthesis and metabolism, thereby affecting the balance between ABA and GA and ultimately controlling wheat SD and germination.

## Discussion

Increasingly complex climatic conditions characterized by alternating heavy rainfall and high temperatures have begun to severely affect wheat production. PHS, which is caused by high humidity or rainfall just prior to harvest, can negatively affect both yield and quality in cereal grains. Therefore, identifying dormancy genes for improving wheat PHS resistance has become a key focus in wheat breeding programs. As one of the largest TF families in plants, MYB TFs are extensively involved in growth, development, metabolism, and stress response (Jin *et al*., 2025; Yang *et al*., 2025; Yang *et al*., 2025). Members of the R2R3-MYB subfamily play crucial roles in responding to abiotic stresses such as salinity, drought, and low temperature (Chen *et al*., 2013; Xiong *et al*., 2014; Zhu *et al*., 2015; Zhou and Tang, 2019). However, in wheat, only TaMyb10-D (a R2R3-MYB TF) has been verified to enhance SD and PHS resistance by directly activating the transcription of *NCED* to promote ABA biosynthesis (Lang *et al*., 2021). It is not yet clear whether other members of the R2R3-MYB TF family are involved in SD regulation.

Here, combined expression, sequence variation, association, and functional analyses confirmed the positive role of *TaMYB83-7B*, encoding a R2R3-MYB TF, in regulating SD. Two T/C mutations in the *TaMYB83-7B* promoter (−907 bp and −1133 bp) were found to be significantly associated with SD and used to develop two diagnostic CAPS markers for distinguishing strong- and weak-dormancy phenotypes. The superior alleles *TaMYB83-7B-907-C* and *TaMYB83-7B-1133-C* were associated with enhanced dormancy. These findings provide valuable targets and molecular markers for the genetic improvement of PHS resistance in wheat through molecular design breeding.

In addition, phylogenetic analysis revealed that TaMYB83-7B exhibits high homology with the R2R3-MYB family member AtMYB20 in *A. thaliana* (Cao *et al*., 2014). Previous studies report that AtMYB20 functions as a negative regulator of the drought stress response by modulating ABA signaling and stomatal closure (Third *et al*., 2017). ABA is well known to induce and maintain SD, as well as to aid drought resistance. Although wheat and *A. thaliana* are evolutionarily distant, both species share conserved stress response regulation mechanisms (Liu *et al*., 2025; Gardiner *et al*., 2015; Toora *et al*., 2024). Therefore, we speculate that TaMYB83-7B may enhance drought resistance in wheat similarly to AtMYB20 in *A. thaliana*, although this hypothesis currently lacks experimental validation.

The antagonistic balance between ABA and GA determines the transition between dormancy and germination (Yang *et al*., 2025). For example, NtMYB330 mediates SD and germination in tobacco by modulating ABA-GA crosstalk through transcriptional regulation of ABA- and GA-related genes (Zhao *et al*., 2022), whereas ZmMYB59 negatively regulates seed germination by inhibiting GA biosynthesis while promoting ABA accumulation (Zhai *et al*., 2020). These examples suggest that R2R3-MYB TFs regulate SD and germination by influencing the ABA and GA pathways. We observed that transgenic wheat seeds overexpressing *TaMYB83-7B* exhibited significantly increased ABA levels and significantly decreased GA levels, resulting in an elevated ABA/GA ratio. This hormonal shift ultimately enhanced SD and inhibited germination. Further analysis revealed that *TaMYB83-7B* regulated the expression of genes involved in ABA and GA biosynthesis and metabolism, thereby modulating the balance the two hormones. This regulatory pattern aligns with the classical model of SD in cereals, in which the dynamic ABA/GA ratio is the key determinant of dormancy depth and germination timing (Loades *et al*., 2023).

## Conclusions

In this study, we combined expression and sequence variation analyses, molecular marker development, and OE, EMS mutant, and heterologous transformation experiments to elucidate the biological function of the R2R3-MYB family TF TaMYB83-7B in wheat SD. The T/C mutations at −907 bp and −1133 bp in the *TaMYB83-7B* promoter were verified to be associated with SD. Furthermore, *TaMYB83-7B* was found to modulate SD by affecting the balance between ABA and GA. Our findings provide valuable genetic resources and molecular markers for breeding wheat varieties with high PHS resistance.

## Author contributions

QSZ, SJC and LTZ conceived and designed this study. QSZ performed the main experiments and wrote the manuscript. SJC, LTZ, and HFW helped with experimental verification and data processing. WZL, ZWW, GJZ, WBL, CXH, WG, and CC assisted in the training and management of experimental materials and data management. CXM, CC, and HPZ designed the study. HPZ supervised this study and revised the manuscript. All authors read and approved of its content.

